# Identification of disease-specific alleles and gene duplications from 1,600 *Haemophilus influenzae* genomes using predicted protein analyses from an unsupervised language model and clinical metadata

**DOI:** 10.64898/2026.03.12.711436

**Authors:** Phillip R. Palmer, Joshua P. Earl, Joshua C. Mell, Kelvin L. Koser, Jocelyn Hammond, Rachel L. Ehrlich, Sergey V Balashov, Azad Ahmed, Steven Lang, Kevin Raible, Angelina L Wong, Brian Wigdahl, Ravinder Kaur, Michael E. Pichichero, Will Dampier, Garth D. Ehrlich

## Abstract

*Haemophilus influenzae,* a Gram-negative bacterium that is an obligate commensal of the human nasopharynx, is associated with both acute and chronic infections of the ears, adenoids, sinuses, and lungs, as well as pelvic inflammatory disease, sepsis, and meningitis. This diverse array of clinical disease phenotypes is due to *H. influenzae’s* natural competence providing for frequent distributed gene exchange among strains during polyclonal colonizations. We performed whole genome sequencing (WGS) on ∼1000 *H. influenzae* strains and combined these data with ∼600 publicly available genomes. Clinical metadata including isolation site and disease were available for all strains. Our goal was to identify variations in protein sequences inferred from the WGS that correlate with the clinical metadata. We employed the alpha-fold unsupervised machine-learning model to all protein sequences. For each unique AA sequence, a numerical vector was generated reflecting the protein’s biochemical properties. Multidimensional clustering was used to group variants into clusters by distance. For gene groups with two or more clusters, clinical metadata from the strain of origin was overlayed and subsequently tested for correlation with the clustering. This allowed identification of multiple gene groups with clusters of variants that correlated significantly with the patients’ disease. COG category analysis showed that many of the significant gene groups were associated with antibiotic targets. The *TbpA* gene group type was found to be significantly correlated with disease type with three of five variant clusters having a 95% prevalence among strains that were isolated from the lungs of COPD patients or other lower pulmonary tract infections.

## Introduction

*Haemophilus influenzae* is a Gram-negative bacterium that exists as an obligate commensal of the human nasopharynx and upper respiratory tract but can transition to an opportunistic pathogen following viral infection or trauma to the contiguous mucosal surfaces (Ehrlich et al 2008; Duell et al 2016). Strains are classified as typeable, if they have a polysaccharide capsule, or nontypeable if they do not have a capsule. There are six serologically identifiable capsular types referred to as a-f. (Khattak et al 2003). *H. influenzae* type b (Hib) was the primary cause of pediatric meningitis and epiglottitis prior to the development of a vaccine in the late 1980’s against its capsule (Wen et al 2020). In the post Hib vaccine era, there has been a highly statistically significant decline in Hib infections and its recovery from patients (Tsang et al 2008; Soeters et al 2018 & 2019; Zhang et al 202;). However, recently there has been an emergence of both invasive a and e serotypes (Whyte et al 2020; Shuel et al 2021; Topas et al 2022; Ulanova et al 2024;), as well as an increase in invasive non-typeable *H. influenzae* (NTHi) (Agrawal et al 2011; Takla et al 2020). The NTHi remain a major pathogen in pediatric cases of acute and chronic infections including otitis media; adenoiditis; conjunctivitis; and sinusitis (Post et al 1995; Rayner et al 1998; Ehrlich et al 2002; Shen et al 2003; Hall-Stoodley et al 2006; Nistico et al 2011; Hu et al 2021;). In adults, the NTHi are responsible for chronic infections in patients with compromised lung function due to COPD (Murphy et al 2004; Mell et al 2018; Short et al 2021;), and cystic fibrosis (Starner et al 2006), and are also associated with pelvic inflammatory disease (Martin et al 2013)

Many factors are known to contribute to NTHi’s pathogenicity including: immune deception and evasion (Heise et al 2018; Kress-Bennett et al 2016); biofilm formation and structure (Post et al 1995; Rayner et al 1998; Ehrlich et al 2002; Shen et al 2003; Hall-Stoodley et al 2006; Nistico et al 2011; Ehrlich et al 2004; Post et al 2004; Schaudinn et al 2007; Kerschner et al 2010; Devaraj et al 2018; Buzzo et al 2021); antibiotic resistance (Jansen et al 2006; Asbell and DCory 201;); high molecular weight adhesins (Atack et al 2020; Fernandez-Calvet et al 2021; Murphy et al 2023); iron/heme uptake mechanisms (Gray-Owen et al 1995;Morton et al 2009); toxin-antitoxin systems (Daines et al 2004; Ren et al 2012); pH tolerance (Ishak et al 2014) and more. Typically, such factors have been discovered in studies utilizing small sample sizes with well-defined conditions specifically designed to uncover one mechanism at a time.

In recent years it has become possible to search broadly for virulence mechanisms of interest whereby large genomic and meta-omic datasets across variable conditions are used to uncover larger trends associated with pathogenicity that can then be subsequently confirmed in mechanistic studies. These studies were predicated on the realization that an individual *H. influenzae* genome contains roughly 1.8 million base pairs and ∼1,750-1800 genes (Fleischman et al 1995; Harrison et al 2005), but that this represents only a small fraction of the species’ genomic complement. Beginning with the rubric of bacterial plurality (Ehrlich et al 2005), which was based on the highly extensive disease variation displayed by various NTHi clinical isolates in controlled animal model experiments (Buchinsky et al 2007) and the distributed genome hypothesis (Ehrlich et al 2010; Nistico et al 2014; Hammond et al 2020; Innamorati et al 2020), which predicted the pan(supra)genome and led to the development and performance of large-scale whole genome sequencing projects in which the NTHi core and distributed genomes were characterized (Shen et al 2005; Gladitz et al 2005; Hogg et al 2007; Hall et al 2010; Boissy et al 2011) at ∼ 1350 and 3250 gene, respectively. These studies showed that the core genome was a small fraction (< 30%) of the pangenome, but that even among the core genes there was very extensive allelic heterogeneity with SNP rates more than order of magnitude higher than among vertebrate species on average.

Once generated, it became possible to use these large omic data sets for the performance of statistical genetic studies (Eutsey et al 2013; Kress-Bennett et al 2016) and the characterization of repeated gene gains and losses among clonal lineages to identify and the characterize distributed genes, including those that were completely unannotated, that were associated with particular tissue tropisms and disease conditions (Kress-Bennett et al 2016; Moleres et al 2018; Kosar 2023). Allelic variations within clusters of orthologous genes (COGs) have yet to be explored within these NTHi datasets. However, a study in *Escherichia coli* provides support for linking variation in core metabolic genes to antibiotic tolerance (Lopatkin et al 2021)

While gene-based association studies aim to utilize a wealth of information, the overwhelming amount of data can make it difficult to search for significant contributors to disease. Machine learning models represent a way to simplify complex data through the use of artificial intelligence, particularly Transformer models [Vaswani et al 2023)] Recently, an unsupervised learning language model, the Evolutionary-Scale Model version 2 (ESM-2), was developed to characterize proteins (Rives et al 2021). Using the same strategy as language models, the network is trained using masked learning; a process in which some of the sequence is hidden and subsequently predicted. The model weights are updated to improve the prediction through gradient descent [Wolf et al 2020]. In this way, the model develops an internal vector representation of provided sequences, termed an *embedding*, in which similar proteins have small distances between their vectors while dissimilar proteins have large distances (Rives et al 2021). While these embedded vectors cannot be translated directly, they are useful inputs into downstream machine learning techniques as they have useful properties such as: a constant shape, normally distributed outputs, and encode information about the protein. These vectors have been used to predict a wide range of outcomes such as: gene expression (Zhang et al 2022), Cas9 off-target sites (Du et al 2025), HIV co-receptor utilization (Dampier et al 2022) and now entire bacterial genomes (Wiatrak et al 2025) In this work, we applied ESM embedded vectors for all genes and variants from ∼1,600 isolates of *Haemophilus influenzae* to explore variations within each gene group and look for correlations with clinical metadata.

## Results

### Characteristics of *Haemophilus influenzae* clinical isolates collection

Of the ∼1,600 *H. influenzae* genomes used in this study, ∼ 1/3 were obtained from online databases with the remainder sequenced in our laboratory, the majority of which were obtained from the biorepository at the Center for Infectious Diseases and Immunology at Rochester General Hospital Research Institute, maintained by two us (MEP and RK). The clinical phenotypes explored in this study were: health state (sick or healthy), anatomical infection site, and patient age. Our isolates are primarily from patients (70%) vs. healthy subjects (30%) (**Fig 1A**); infection type was by anatomic site to probe for tissue-specific adaptations (**Fig 1B**). We also parsed for age: senior > 65 years; adult 18-65 years; child 2-18 years; and baby < 2 years of age (**Fig 1C**). Gene groups (GG) were defined from the pan-genomic analyses as protein sequences containing 75% or greater homology (Hogg et al 2007) resulting in 4,610 GG for analysis.

**Figure 1.**
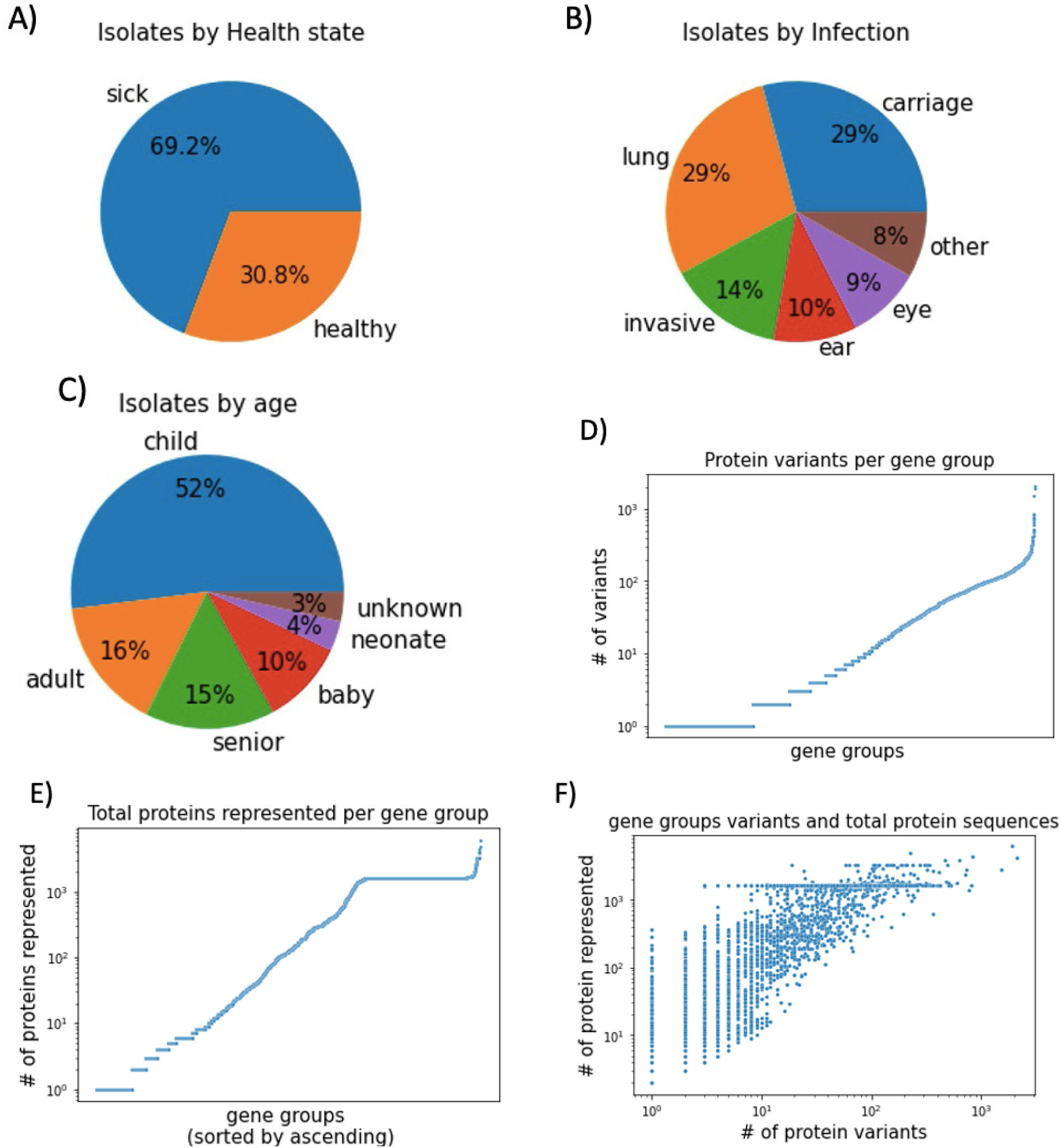
Summary of *Haemophilus influenzae* clinical isolates and their genes. Percentage of all isolates by: health state (A), infection type (B), and age (C). Gene groups of protein sequences containing >75% similarity were analyzed. Shown are total number of sequence variants per gene group (D), and total number of protein sequences (redundant sequences included) per gene group (E), sorted by ascending. These values were then plotted together as unique variants vs total protein sequences (F).

Within these gene groups are variants; represented by each unique protein sequence. Total variants for each gene group were plotted in ascending order (**Fig 1D**). Many gene groups contained a single variant, and about half of all gene groups have less than 10 variants. At the other end of the spectrum, a few gene groups contained many variants. Low variant total is consistent with such genes being highly conserved, as expected with core genes. A single sequence variant can represent any number of strains that contain that sequence. Therefore, total proteins represented were enumerated for each gene group and also plotted in ascending order (**Fig 1E**). A large number of gene groups contain roughly 1,600 total sequences, indicating roughly one per strain (1,600 total strains). Any gene groups with a value greater than 1,600 contain, on average, more than one copy per genome. However, when the two data sets were combined as total protein sequences represented versus number of variants, the gene groups with roughly 1,600 total sequences showed a large amount of sequence diversity (**Fig 1F**). Surprisingly, there were no gene groups with only one variant that were present less than 600 times, suggesting that there is a large amount of genotypic diversity amongst all isolates, even in core genes.

### Process overview of machine learning model application and analysis

A pipeline was constructed to explore intragenic diversity of *H. influenzae* for hidden properties that correlate with various clinical phenotypes (**Fig 2A**). The crux of this algorithm is a previously developed unsupervised machine learning language model (Rives et al 2021). This model uses only amino acid sequence information which is then converted to a numerical vector that reflects biological aspects of the protein. This is done by inferring meaning of the amino acid from its context within the protein, similar to words in a sentence.

**Figure 2.**
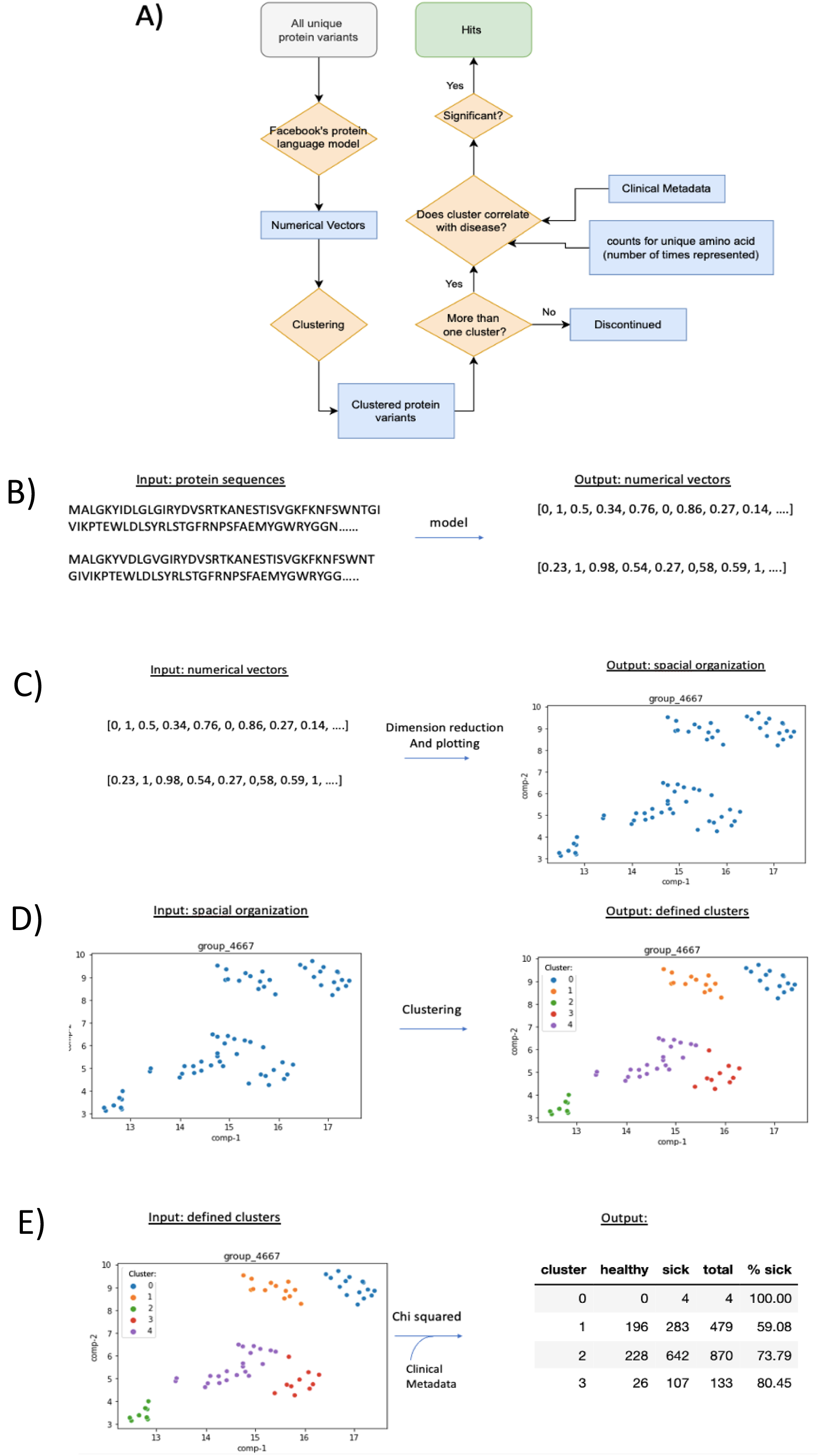
Process overview and sample input and outputs. A) Flowchart of the application and analysis of an unsupervised language model on protein sequences to identify clusters of variants that correlate with clinical metadata. B) Modeling: protein sequences are processed by the model and converted to numerical vectors. C) Visualization: numerical vectors are shown in reduced dimensions to visualize proximity. This step was not included in the main pipeline; it is shown here for visualization purposes only. D) Clustering: HDBScan was used to generate clusters from multidimensional positioning E) Correlation with metadata: Chi squared analysis was performed to check for correlations between cluster and clinical metadata

An example of this transformation is shown in **Fig 2B**. These numerical vectors are bulky and can be simplified for visualization by dimension reduction, however this step was only done in this study for visualization and was not included in the main pipeline to maintain reproduceable results. Clustering occurred at the multidimensional level, prior to dimension reduction. We used uniform manifold approximation and projection (UMAP) to reduce the vector to two dimensions so that it can be plotted for visualization (**Fig 2C**). Each point represents a unique protein sequence within the gene group. Clustering was performed using HDBScan with a minimum threshold of six variants for a cluster (**Fig 2D**). Variants identified as not belonging to a cluster (noise) were dropped. Because we were interested in comparing multiple clusters within one gene group, gene groups with only one cluster were discontinued. A gene group with two or more clusters then had additional data for total protein counts appended. This is because a single sequence variant often represents many occurrences of that sequence. Clinical metadata was also retrieved and appended to the protein data table, corresponding to the strain harboring each protein. A contingency table of cluster and the category of interest (ex: infection type) was created and a Chi squared test of independence was calculated. A p-value of 4×10^-5^ was determined to be statistically significant based on a Bonferroni correction corresponding to the number of gene groups that were analyzed (groups containing more than one cluster). The resulting significant genes are reported herein, and further analysis was conducted depending on the category of interest.

### Correlations of cluster and health state

To evaluate the machine learning model’s ability to find meaningful differences amongst variants, we looked for correlations between variant cluster and health state. Health state here refers to isolates from healthy or sick patients. Isolates with an unknown health state were removed from the analysis. For gene groups with two or more clusters, a contingency table was generated for cluster and health state to enumerate counts for each combination of cluster number and health state. To determine the relationship between cluster and health state, a Chi squared test of independence was performed. If the p-value was less than 4×10^-5^ the gene group was considered significant, as discussed in the previous section. 79 gene groups were found to be significant for dependence of cluster and heath state (Table 3.1). Of those significant gene groups, nine had at least one cluster with greater than 95% isolates from sick patients, and five of those had a cluster that was 99% or higher (**Fig 3A**). Gene group hcat had less than 1% sick. This analysis identified clusters enriched for a particular health state. To identify which cellular processes were represented by the gene groups that were significantly associated with one of the test states, COG categories were predicted by functional domain analysis using eggNOG-mapper (Cantalapiedra et al 2021) to predict. The most prevalent amino acid sequence for each gene group was used for this analysis. Unknown function was the most represented COG category (**Fig 3B**), although it was also the most numerous category amongst all of the gene groups. When normalized for change in prevalence, defense mechanisms and inorganic ion transport categories showed the largest change (**Fig 3C**). A contingency table for gene group tbp1 shows clusters containing predominantly sick isolates (**Fig 4A**). tbp1 was no longer significant when strain duplicates were removed from each cluster, yet it remains of interest because of its high percentage of sick isolates in many clusters (**Fig 4B**).

**Figure 3.**
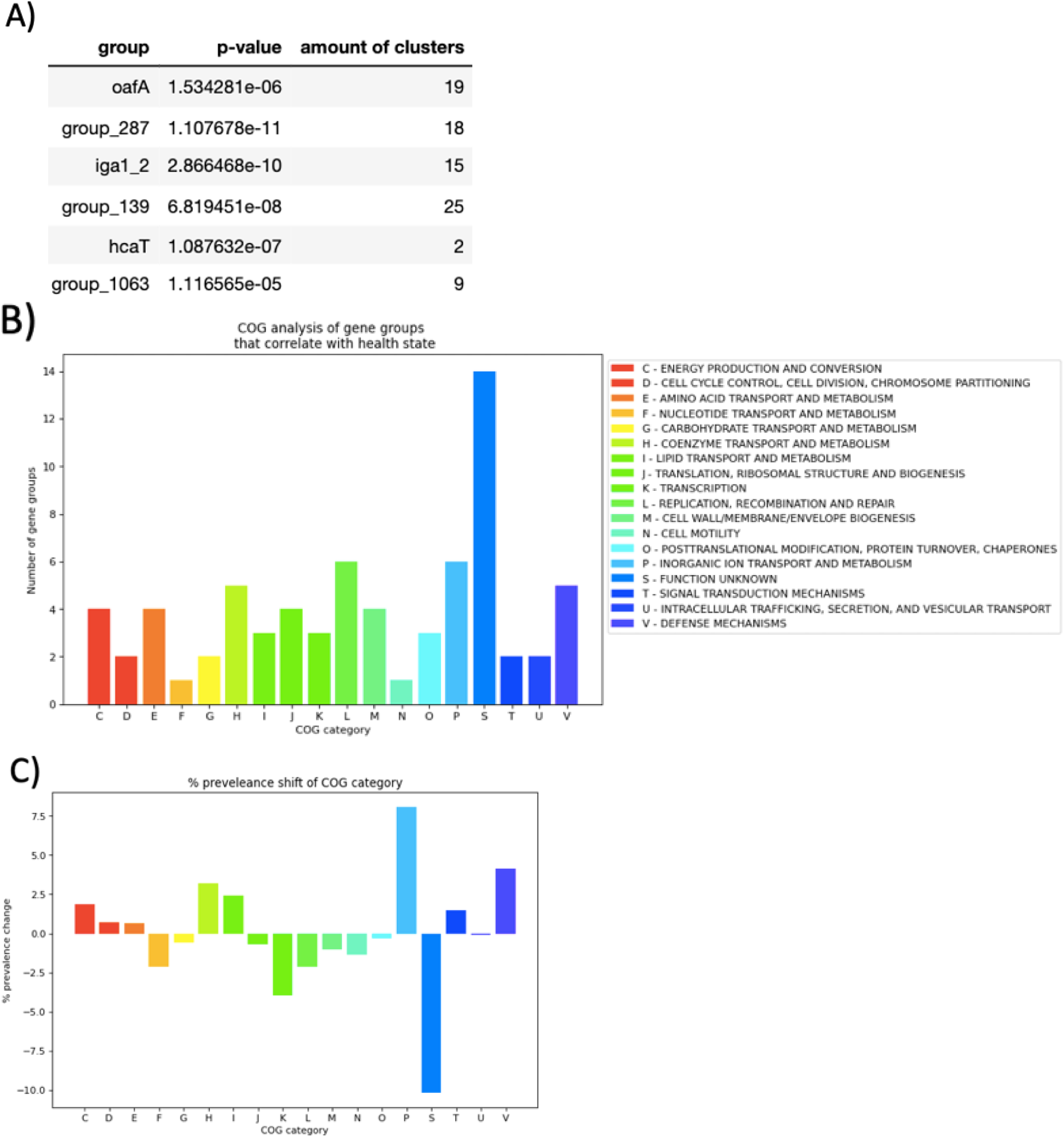
Correlations of cluster and health state. A) Significant gene groups that had at least one cluster with >99% sick or healthy isolates. B) COG category analysis of all significant gene groups. C) Change in prevalence of each COG category.

**Figure 4.**
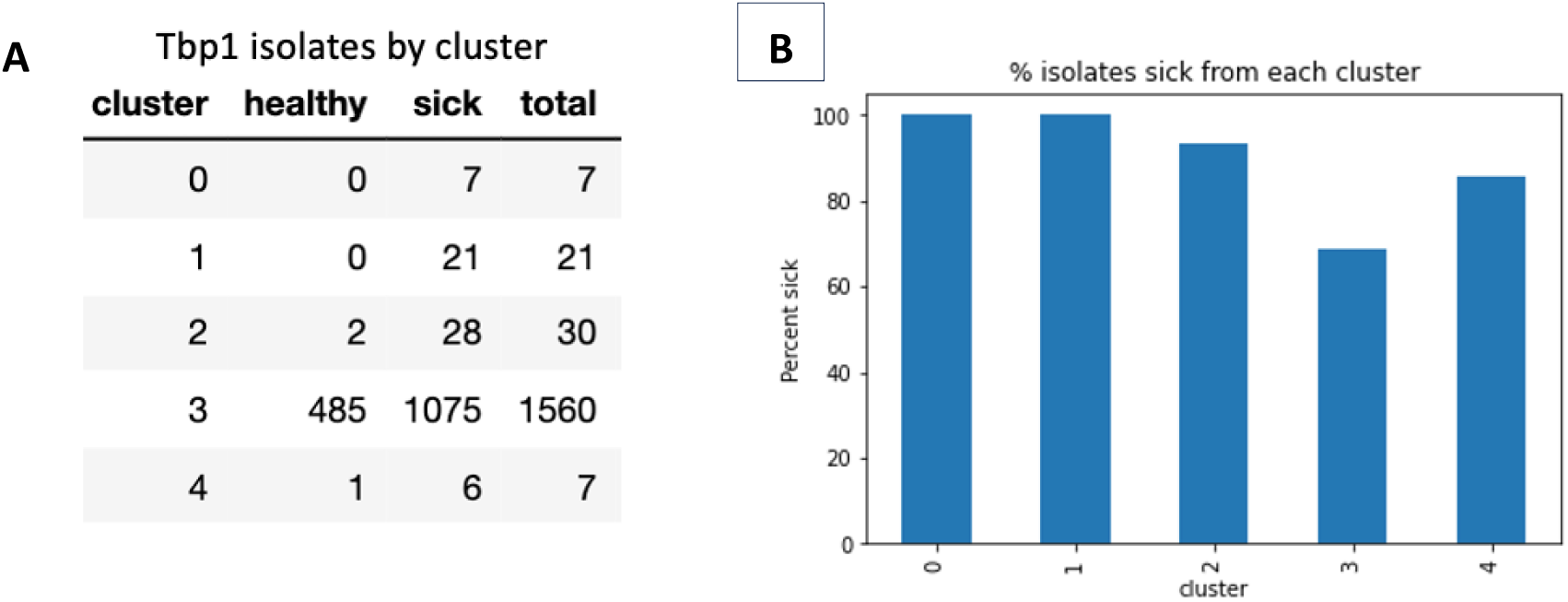
A) Contingency table of cluster and health state for gene group *tbpA*. B Plotting *tbpA* clusters by percentage of isolates that are sick.

### Correlations between clusters and infection type

All identified clusters were then overlaid with the tissue/organ (ear, lung, carriage, eye, invasive and other) from which the corresponding isolate was recovered, and a contingency table was created of number of clusters versus infection type. This was used to conduct a Chi squared test of independence. Using the same p-value as above, 285 gene groups were found to be significantly associated with the tissue of origin (**Table 2**). Four of the gene groups with significant associations contained a cluster with ³ 99% prevalence for one or more origin sites, ten from the lung and two NP carriage (**Figure 5A**). Three contained a cluster with ³ 99% prevalence in the lung, and the *hcat* gene group was predominantly associated with carriage. COG category analysis of significant gene groups showed many with: unknown function; cell wall synthesis; and amino acid transport and metabolism (**Figure 5B**). After normalization to the greater gene group collection, only the latter two showed an increased in prevalence (**Figure 5C**), both of which are common targets of antibiotics. Tbp1 was a top hit again; two clusters had a 100% prevalence in the lung, and a third had 93%. Analysis of the contingency table for *tbpA* clusters versus infection type or health state, it was observed that four clusters from *tbpA* are predominantly associated with isolates from the diseased lung, while the remaining cluster resembled the greater collection (**Fig 6A&B**)

**Figure 5.**
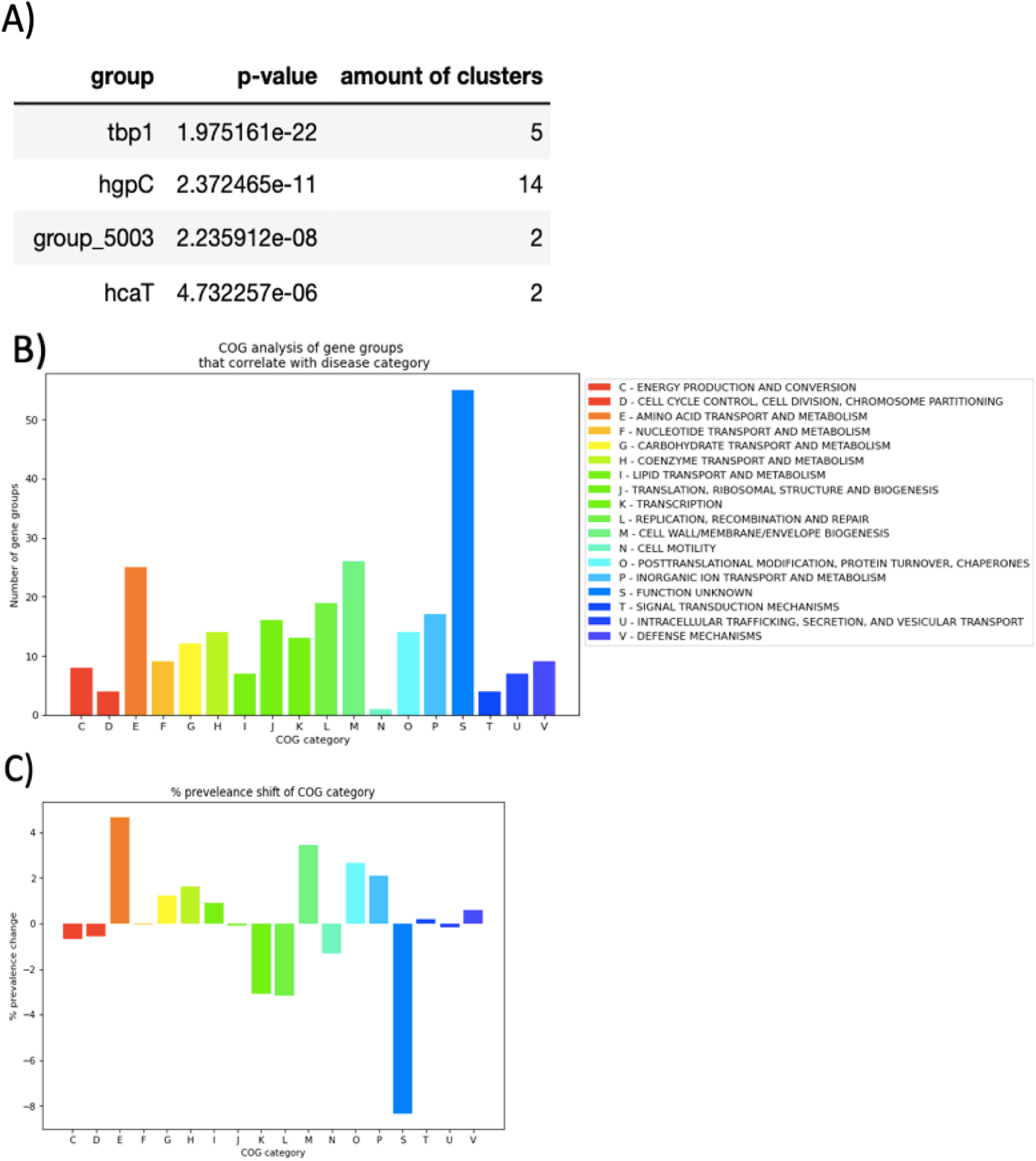
Correlations of cluster and infection type. A) Significant gene groups that had at least one cluster with >99% one infection type. B) COG category analysis of all significant gene groups. C) Change in prevalence of each

**Figure 6.**
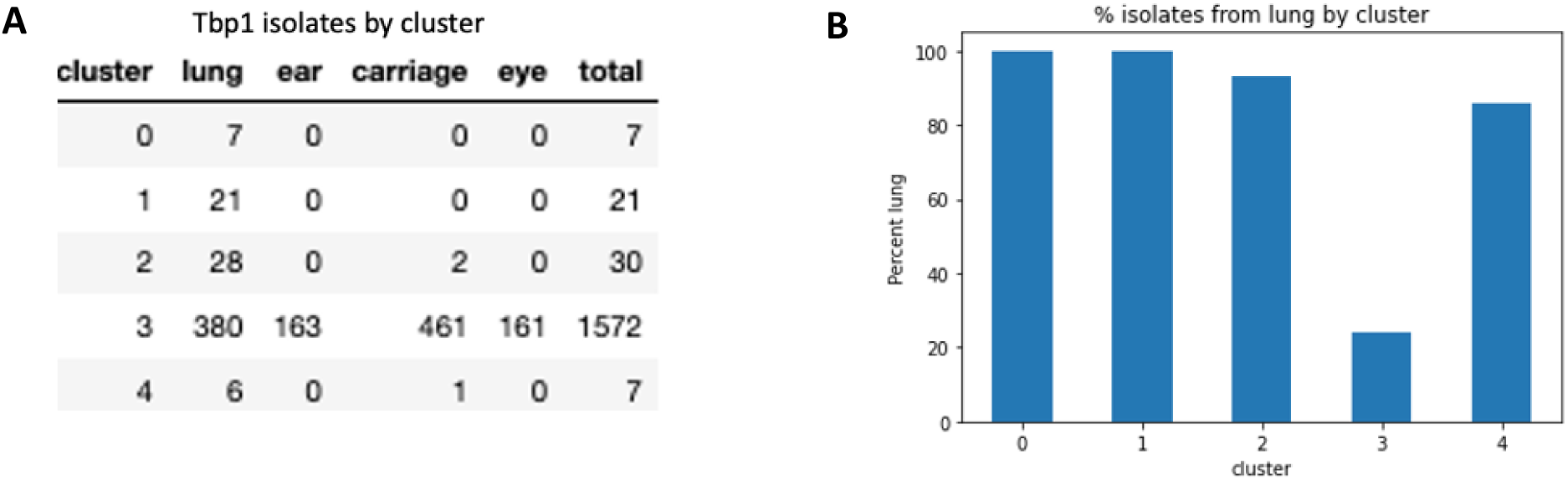
Correlations of cluster and infection type. A) Contingency table of *tbpA* cluster and infection type. B) Plotting *tbpA* clusters by percentage found in the lung.

### Analysis of *tbpA*

*tbpA* was selected for further exploration because four of its five clusters contained a higher percentage of isolates that were from lung infections. *tbpA* encodes an outer membrane transferrin binding protein, Tbp1; that serves as a virulence factor involved in iron uptake from the host. It is essential for *H. influenzae* growth on media with human transferrin as the sole iron source (Gray-Owen et al 1995). Our analysis showed that gene group *tpbA* consists of 421 unique variants from 1,782 total protein sequences. Modeling and clustering of the unique amino acid sequences resulted in five clusters. Cluster 3 includes 368 protein variants and 1596 total proteins, and in terms of infection type prevalence, resembles the greater collection. Clusters 1 and 2 represent 21 and 30 strains respectively, all of which were isolated from patients with lung illnesses. Clusters 4 and 0 are also primarily found in isolates from the lung, but each of these clusters consist of < 10 strains. Reducing dimensions of the numerical vectors shows the spatial proximity of the clusters in 2 dimensions, down from 1,280 (**Fig 7A**). Some differences among the clusters show low resolution due to clustering prior to dimension reduction. However, the overall image shows a general separation of clusters 0, 1, 2, and 4 from cluster 3. Clusters 0, 1, 2, and 4 displayed a reduced average protein length compared to cluster 3 (**Fig 7B**) suggesting a truncation of the gene. A pfam domain analysis of the variants in each cluster revealed a significant decrease in the prevalence of either or both of the receptor and plug domains, from the isolates recovered from lung patients (**Fig 7C**). We noted that all strains present in clusters 0, 1, 2, and 4 were also present in cluster 3. Therefore, it is likely that these clusters are truncated duplications that occurred in strains that have a full-length copy of *tbpA*. Due to the small number of isolates found in these clusters we sought to determine if this phenomenon was confined to (possibly related) strains obtained from a single collaborator, but this was not the case as these clusters consisted of isolates from two unrelated cohorts: cystic fibrosis isolates from Seattle, Washington USA, and COPD isolates from Madrid Spain. This observation suggests that this phenomenon of probable *tbpA* partial duplication has occurred repeatedly within patients with different lung diseases

**Figure 7.**
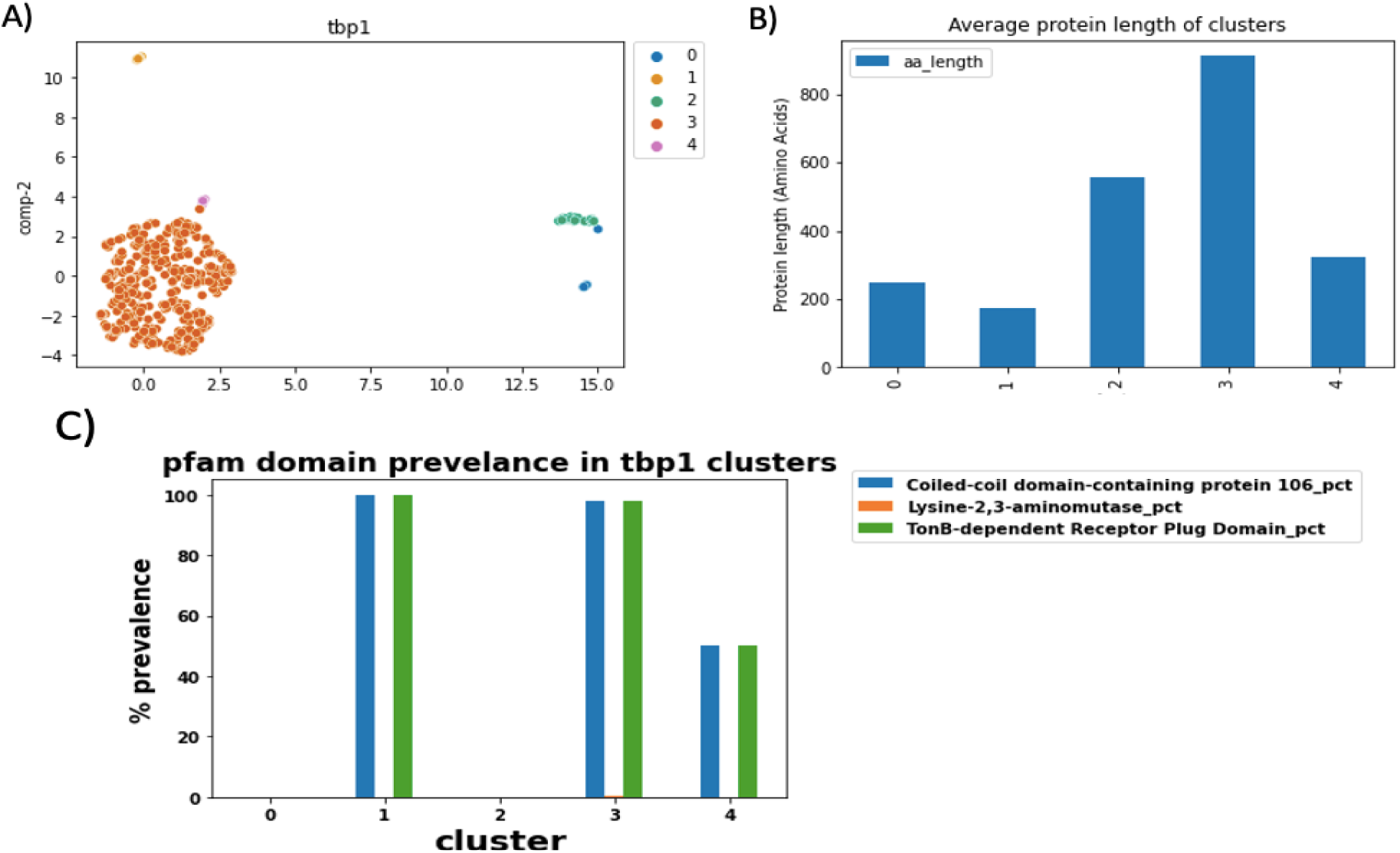
Analysis of Tbp1. A) clustering of Tbp1 sequence variants as classified by the model. B) Average amino acid length of Tbp1 by cluster. C) Prevalence of Pfam domains by cluster

### Correlations of cluster and patient age

Finally, we correlated variant clusters with patient age. 286 gene groups were found to be significant by a Chi squared analysis between cluster number and age group (Table 3). Of the significant gene groups, only three had at least one cluster with greater than 99% prevalence of isolates belonging to a single age group (**Fig 8A**). These were annotated as: *hgpB*, a hemoglobin-haptoglobin binding protein; group_287, a predicted glycosyltransferase family 8 protein; and *msfA_1*, a hypothetical related to a previously characterized virulence factor by our group (Kress-Bennett et al 2016). A COG category analysis revealed that the significant gene groups are enriched in unknown function, amino acid transport and metabolism, translation, and cell wall synthesis (**Fig 8B**). After normalization, amino acid transport and metabolism, inorganic ion transport and metabolism showed the largest increase in prevalence (**Fig 8C**).

**Figure 8.**
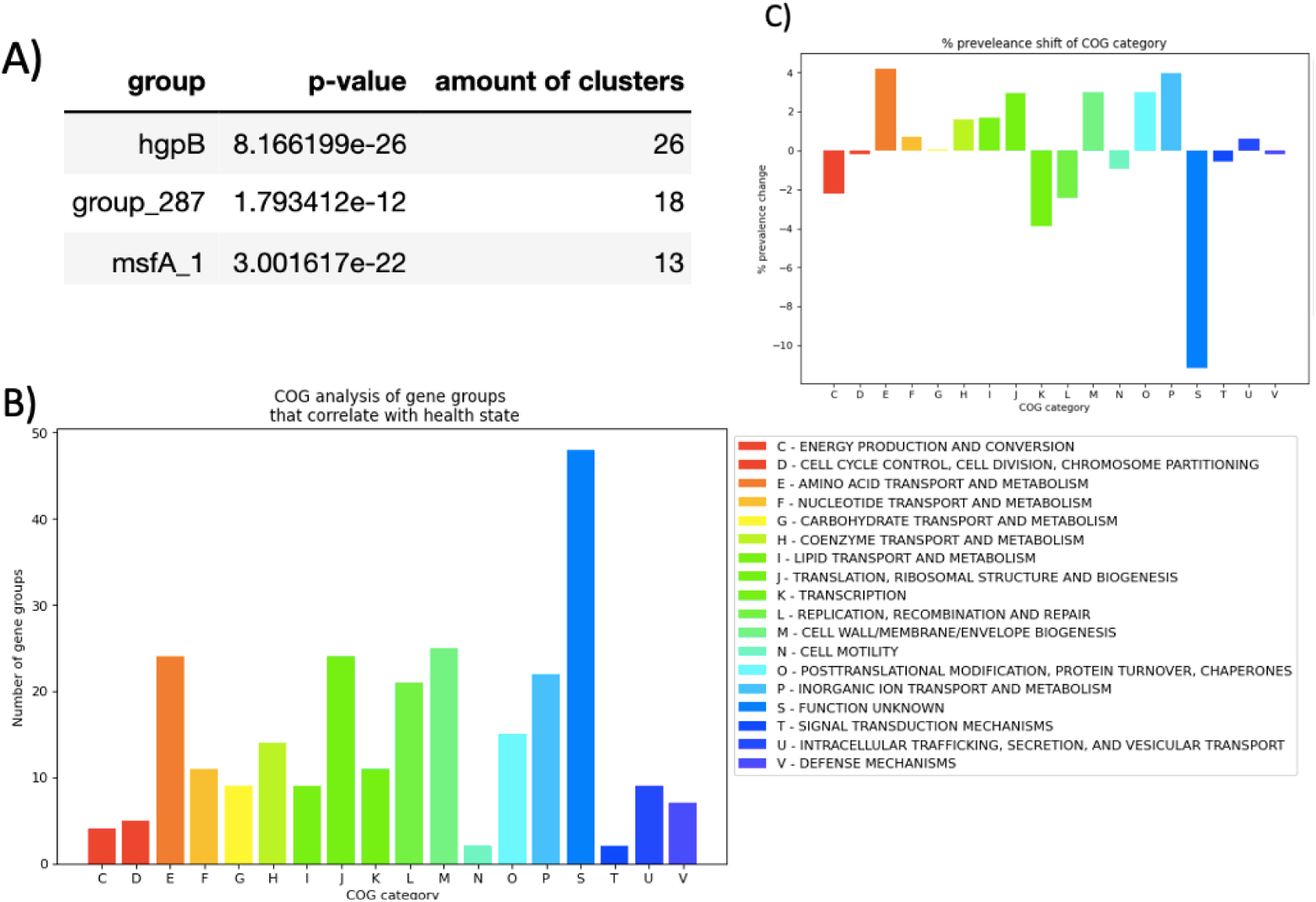
Correlations of clusters and patient age. A) Significant gene groups that had at least one cluster with >99% in one age group. B) COG category analysis of all significant gene groups. C) Change in prevalence of each COG category.

## Discussion

In this study, we developed a method for studying clinical implications of protein variants by implementing a previously described machine learning model. ∼1,600 clinical isolates were analyzed using this method to look for correlations between protein variants and clinical metadata including as health state, site of infection, and patient/subject age.

70% of all isolates evaluated were from diseased individuals, which could affect results when searching for clusters enriched for sick isolates. Because the expected ratio of sick isolates will already be high, this data may be biased compared to a dataset of equal parts sick and healthy isolates. It is important to note that the pathogenicity of an isolate isn’t determined by a patient’s health state; it is possible to pull a commensal isolate from a sick patient, and a pathogen from a healthy patient. Additionally, some isolates were serial isolates from the same patients over a time course, which may lead to over representation for genes from one lineage of isolates.

This analysis pipeline is reproducible and uses open-source tools that are listed in the methods section. It is important to note that other tools used for the same purpose, or more recent versions of tools used herein may provide variations in results. The data used in this study is available as part of a concurrent publication from our lab in which the dataset is described (Hammond et al submitted – see appendix).

Because the pangenomic tool Roary was used to create gene groups, they often have arbitrary “group_###” names. However, some groups retained a common gene name. Of the 4,610 gene groups observed, roughly 70-300 gene groups per category were found to significantly correlate with clinical metadata following a Chi squared contingency test. These hits will serve as preliminary data for future studies investigating the role of allelic adaption of pathogens to specific organs and disease states. Three separate analyses were conducted for correlations of variant clusters with health state, infection type, and patient age. Health state (sick or healthy) resulted in top gene groups with some clusters containing greater than 99% sick derived isolates such as the O antigen gene oafA. The O antigen is a key antigen in host recognition, providing a potential rationale for the oafA gene groups correlation with a sick state of health. Additionally, other virulence factors were found amongst top hits including cell adhesion, iron acquisition, and other outer membrane proteins (Table 3.1). This is promising evidence that this pipeline can be used successfully to identify the evolution of novel virulence factors associated with specific disease states.

COG category analysis of gene clusters significantly associated with specific conditions (for all three analyses) showed results for inorganic ion uptake, common targets of antibiotics such as cell wall biogenesis and translation. The finding of COGs associated with amino acid transport and metabolism are of note due to the stringent response’s ability to upregulate persistence in response to amino acid starvation (Traxler et al 2008). Similar results were observed for cluster versus infection type.

*tbpA* was a top hit associated with an infection type, as three of its five clusters had over 94% of isolates that were recovered from the lungs of CF and COPD patients. We focused on its gene product Tbp1, a transferrin binding protein, because iron uptake is a known virulence factor and is essential for *H. influenzae’s* survival in the host. It was observed that four of the *tbpA* clusters appeared to be truncated duplications, perhaps as an attempt to increase iron uptake from the host. Interestingly *tbpA* is a known hotspot for DNA recombination in *H. influenzae* (Yahara et al 2016).

### Conclusions and future directions

We have established a pipeline for classifying proteins into intragenic clusters and analyzed those clusters for correlations with clinical metadata. COG category analysis showed that many of the affected pathways are common pathogenic factors or antibiotic targets. In particular, the gene group *tbpA* was identified to have multiple clusters that had a high prevalence of isolates from the lungs of CF and COPD patients. Overall, this pipeline will serve as a resource tool for identifying genes of interest in clinical infections of *H. influenzae* and other pathogens for which there are a sufficiently large enough number of genomes for which there is clinical metadata.

The identification of genes encoding only hypothetical proteins is of particular interest as these findings suggest a rational means to target COGs within the genomic dark matter that are likely associated with some aspect of virulence. Domain and motif analyses that distinguish clusters within a given COG that are associated with a specific aspect of clinical disease can be used to ascertain how such domains are responsible for producing the observed effects.

## Methods

### H. influenzae isolates

Of the 1618 *Haemophilus influenzae* genomes analyzed in this study, 896 were newly obtained from selected clinical isolates representing all major diseases and tissues from which the NTHi are isolated. Isolates were selected from across several different cryo-collection repositories to represent a diverse range of geographical and clinical sources, with a focus on isolates from under-represented categories in the public databases, namely from nasopharyngeal carriage, conjunctivitis, cystic fibrosis lung, and several rarer categories; these included adenoid biopsies in children, endometritis-associated with pelvic inflammatory disorder in women, and others. For each isolate, available information was compiled that, where possible, recorded the original date of isolation, geographical location, subject age, health status, a detailed description of clinical provenance, and categorization into one of six broad classes: carriage, ear, eye, invasive, lung, and other. Most isolates were from distinct subjects, though some were collected from the same subject at the same or different times.

To obtain high-quality genomic DNA (gDNA), cultures of *H. influenzae* were grown following standard procedures (Poje & Redfield 2003; Shen et al 2005; Hogg et al 2007); briefly, minimally passaged clinical isolates were streaked on chocolate agar or sBHI agar (brain-heart infusion supplemented with 2 μg/ml NAD and 10 μg/ml hemin) and incubated overnight at 37°C with 5% CO_2_. Single colonies were inoculated into 5 mL overnight shaking cultures in sBHI broth at 37°C. Genomic DNA was extracted from washed cell pellets using the Qiagen DNeasy® Blood & Tissue Kit following manufacturer’s recommendations, and quality and quantity were evaluated by agarose gel electrophoresis, spectrophotometry (NanoDrop^TM^ 1000, Thermo Scientific^TM^), and fluorometry (Quant-IT dsDNA Assay Kit, Molecular Probes, Life Technologies, New York, NY).

### Genome sequencing and assembly

Whole-genome shotgun sequencing of gDNA was performed using either Illumina or Pacific Biosciences (PacBio) technologies. Illumina libraries were constructed using Nextera XT kits following the manufacturer’s protocols and sequenced on a NextSeq 500 using mid-output kits on pools of 96 barcoded libraries to collect 2×150 nt paired-ends with a targeted depth of 200-fold coverage. PacBio libraries and single-cell real-time (SMRT) sequencing were performed according to manufacturer’s recommendations on an RSII instrument (v2.1.0) using 8 to 12 isolates per SMRTcell with a targeted depth-of-coverage of 100-fold.

The assembly pipeline used depended on the sequencing technology. Illumina data were assembled using SPAdes (v3.7.0) (Bankevich et al 2012: Prjibelski et al 2020), after FastQ generation and demultiplexing were performed with bcl2fastq (v2.17.1.14), technical sequence removal using COPE, and error correction with ErrorCorrectReads.pl from allpathsLG. PacBio data were assembled using HGAP (v2.3) (Chin et al 2013) after initial processing through SMRTanalysis (v2.3). Circlator (v1.0.2) (Hunt et al 2015) was used on initial assemblies with permutation to the *dnaA* gene, followed by error correction on the circular junctions using SMRTanalysis resequencing protocols; uncircularized and multi-contig PacBio assemblies were not subjected to this procedure.

Assemblies or contigs were dropped from further analysis if they failed any of the following quality control steps. Illumina SPAdes assemblies were removed if <60-fold coverage was obtained, and individual contigs were removed when: k-mer coverage was <8, contig length was <256 nucleotides, or sequence composition consisted of >95% from a single base. PacBio assemblies were dropped if <10-fold coverage was obtained, and contigs were filtered based on corrected coverage (sum of all bases in the contig divided by the sum of the coverages from all bases in the assembly) below a specific threshold of 0.2.

### Curation of *Haemophilus influenzae* genome assemblies from publicly available sources

Genome assemblies of 652 clinical strains of *H. influenzae* were downloaded from RefSeq, Sanger Center (De Chiara et al 2014) or provided by the CDC on or before 2018-07-23. Additional assemblies uploaded to RefSeq on or before 2023-05-31 were also downloaded for later analysis. For each assembly, metadata was collected by identifying, where possible, information about strain provenance as above, in addition to literature citations associated with specific isolates, as well as obtaining deanonymized clinical metadata via request from the original submitters where possible (Appendix A).

### Quality control filtering of assemblies

To ensure consistent quality among new and downloaded assemblies, each had to pass these additional quality control criteria: first, assemblies were checked for their taxonomic assignment to *H. influenzae* using taxator-tk (v1.2) (Dröge et al 2015) with the nonredundant-microbial_20140513 database; incorrect or higher taxonomic classifications to *Haemophilus*, *Pasteurellaceae*, or bacteria were not considered further. Assemblies from all were removed, if: (a) they were produced by Illumina SPAdes but had >100 contigs; (b) they were produced by PacBio but had >10 contigs; (c) were scaffold assemblies with >250 ambiguous base pairs; or (d) were deprecated or removed by NCBI due to high pseudogene content (typically found for 454 assemblies). Assemblies were also removed if their total length was less than 1.6 MB or above 2.3 MB, as all *H. influenzae* assemblies on RefSeq were between 1.75 and 2.11 MB at the time of download. Finally, a duplicate filter was applied to remove redundant assemblies derived from the exact same clinical isolate; the assembly with lower contig count was typically retained. In some cases, potential duplicates were retained if the provenance of the strains was not obvious.

### Gene annotation and clustering of homologous protein-coding sequences

To ensure consistency, Prokka (v1.13) was used for gene annotations from all assemblies (Seemann 2014), including re-annotation of all publicly available genomes. Gene naming used a custom database of *Pasteurellaceae* genomes from NCBI (67). *In silico* capsule serotypes were determined using hicap with default settings (v1.0.3) (Watts and Holta 2019). This software annotated samples based on the presence of an intact *cap* locus. The presence of capsule was determined by the presence of regions I, II, and III of the capsule locus, and specific serotypes *a* through *f* were delineated by the variant present in region II of the *cap* locus.

Using the annotated assemblies for each strain, homologous gene clusters were grouped using one of two methods: (a) the rapid large-scale prokaryote pangenome analysis pipeline Roary (v3.12) (Page et al 2015); or (b) Panaroo (v1.3.2) (Tonkin-Hill et al 2020). Both programs identify homologs on the basis of an amino acid identity threshold determined using a procedure equivalent to all-by-all BLASTP. On the basis of previous work (Hogg et al 2007; Ahmed et al 2011; Molares et al 2018), a BLASTP threshold of 75% was used to cluster homologs unless otherwise stated. Panaroo was run using ‘strict’ mode to remove erroneous gene clusters and other potential sources of contamination. For both tools, paralogs were not split apart, unless otherwise indicated. The resulting gene presence-absence matrix provided the dataset for subsequent statistical association and machine-learning classification. Gene presences and absences were visualized using pheatmap in R (v1.0.12) (Kolde 2019).

### Supra-genome analyses and population structure

Codon-aware alignments of all protein-coding genes were made using PRANK (v.100802) (Löytynoja et al 2014). The resulting alignments from genes present in at most a single copy per strain were concatenated and used to generate a species-level phylogeny using RAxML (Random Axelerated Maximum Likelihood) (v8.2.4) (Stamatakis 2014). To root the tree, assemblies of two of the closest-related species, *Haemophilus haemolyticus* and *Haemophilus parainfluenzae*, were downloaded from RefSeq. These 13 *H. haemolyticus* and 33 *H. parainfluenzae* assemblies were annotated using Prokka as described above, then combined with the 1618 *H. influenzae* annotations to create a maximum-likelihood tree using RAxML. The midpoint between the *H. influenzae* and *H. haemolyticus* clades was set as the root for the original tree containing only the 1618 *H. influenzae* strains. Tree visualization used APE (v5.7) and ggtree (v3.2.1) packages in R (Paradis & Schliep 2019; Xu et al 2022)

Identification of specific lineages used two comparable methods: (a) *in silico* multi-locus sequence typing (MLST)(Jolley et al 2004) (v2.23), and (b) Population Partitioning Using Nucleotide K-mers (PopPUNK) using default settings (Lees et al 2019).

### Unsupervised machine learning analysis pipeline

Genes were grouped by 75% sequence similarity using Roary for pangenome analysis (80). All unique protein sequences were processed by the protein language model (Rives et al 2021) into numerical vectors with a length of 1,280 values. Each gene group was then individually analyzed for significance of variants. All unique genes within a gene group were collected and the vectors were clustered using HDBScan version 0.8.28. Unique genes labelled as noise were removed, and further analysis was performed if two or more clusters of unique genes were found. Unique gene data of total protein counts, and clinical metadata were then merged to the clustering data. A contingency table was generated for clinical category versus cluster and a Chi squared analysis was performed to generate a p-value for correlation of clinical status and variant cluster.

### Eggnog annotation and COG category analysis

For annotation of all significant gene groups, the most common protein variant was used. A FASTA file was then submitted to eggnog-mapper (Cantalapiedra et al 2021) with smart annotation (<u>eggnog-mapper.embl.de/</u>). COG category raw counts and prevalence (% hits - % overall) were plotted using Seaborn.

### UMAP analysis of tbp1

Uniform manifold approximation and projection (UMAP) was used to reduce dimensionality for the vectors from roughly 1,200 to two components for plotting and distance between points. For reproducibility, the random state parameter for UMAP was set as: *random state=42*. The resulting 2D array was plotted so that each point represented a unique amino acid sequence.

### Pfam domain analysis with hmm-scan

All unique protein sequences of tbp1 were analyzed using hmm-scan with the Pfam database. Prevalence for each Pfam domain prevalence was calculated as percent of each cluster.

## Acknowledgements

This work was supported by NIH grants DC 02148 and DK 082316 to GDE and R01 MH110360 to BW with additional support from the Oskar Fisher Project, a James Truchard Philanthropy, and the Bill and Marian Cook Foundation to GDE.

## Data Availability

The *Haemophilus influenzae* genomes sequenced and used for the analyses herein are available at the NCBI under BioProject PRJNA450095

